# Environment and physiology shape antiphage system expression

**DOI:** 10.64898/2025.12.14.694197

**Authors:** Lucas Paoli, Baptiste Laruelle, Rachel Lavenir, Arthur Loubat, Florian Tesson, Baptiste Gaborieau, Aude Bernheim

## Abstract

Bacteria and archaea encode on average ten antiphage systems. Quorum sensing, cellular, or transcription factors can regulate specific systems (CRISPR-Cas, CBASS). Yet, a systematic assessment of antiphage systems expression patterns is lacking. Here, we combine publicly available RNA-seq data from 14 different species with an original RNA-seq dataset of 15 *Escherichia coli* strains across six environmental conditions and two growth stages. Using this data, we explore the transcription patterns of 236 antiphage systems from 81 types. Defense system expression is variable along environmental, physiological, as well as spatial gradients, and can correlate with cellular physiology and mobile genetic element activity. We identify antiphage systems as cohesive but complex transcriptional units, find coordinated expression of defense islands possibly underpinned by local regulators, and demonstrate the functional relevance of differential expression in native systems. Together, these results suggest that environmental and physiological factors regulate prokaryotic immunity and may prime bacteria for infection.

---

Bacteria and archaea harbour a vast repertoire of antiphage systems to resist infection by virus and other mobile genetic elements. With the recent discovery of more than 200 antiphage defense systems^1^, prokaryotic genomes are now thought to encode an average of at least ten such defenses^2^. A number of these prokaryotic antiphage defenses are evolutionarily related to eukaryotic immune proteins^3–5^. In eukaryotes, the immune response is largely driven by transcriptomic upregulation of immune genes following pathogen detection^6^. By contrast, the extent, drivers, and nature of transcriptional regulation in prokaryotic immunity remains largely unexplored. This limitation stems in part from the fact that most antiphage systems have been characterised through plasmid-based heterologous expression rather than in their native genomic context^7^.

Phage infection was shown to induce transcriptional upregulation of defenses in a few specific cases^8–10^, but the prevalence of this response is uncertain. The phage lytic cycle can be extremely rapid (sometimes under 20 minutes^11^) and many phages degrade host genomes^12,13^ limiting the possibility of mounting a transcriptomic response to infection. By contrast, studies on the regulation of antiphage systems such as CRISPR–Cas or CBASS have shown that their expression is largely driven by phage-independent cues, including quorum sensing and metabolic status^14–22^. Beyond these specific examples, we currently lack a broad assessment of defense system transcription levels, their variation across physiological and environmental gradients, as well as across strains or species. Shedding light on these transcriptional patterns will however be key to understanding the regulation of prokaryotic immunity and its relevance in ecological settings, as well as untangle the potential synergies, redundancies, and fitness costs associated with encoding numerous antiphage systems per genome.

Here, we combine an original, highly resolved RNA-seq dataset of 540 samples spanning 15 *Escherichia coli* strains, six environmental conditions, and two growth stages with around 10,000 publicly available RNA-seq samples from 15 different species^23^ to explore the transcription patterns of 234 antiphage systems across 78 subtypes. Our analyses provide a systematic assessment of defense system transcription levels, variability, and drivers. We further unwind the genetic basis of transcriptional regulation and demonstrate how this information can guide the validation and study of prokaryotic immunity in native settings.

## Results

In order to explore the expression of antiphage systems across conditions, species, and strains, we aimed to associate a large-scale database spanning distant species with a strain-resolved dataset. To that end, we combined the iModulons database, around 10,000 publicly available RNA-seq samples from 14 prokaryotic species (figure 1A, Methods)^23^, with the Picard-T dataset: an original, highly resolved *Escherichia coli* RNA-seq dataset spanning 15 strains (Figure 1E). The iModulons database contains a reference strain for model species from major bacterial lineages, including Actinomycetes (*Streptomyces albidoflavus* and *Mycobacterium tuberculosis*), Gammaproteobacteria (*Pseudomonas* spp., *Escherichia coli*, *Salmonella enterica*, *Acinetobacter baumannii*, and *Vibrio natriegens*), Bacilli (*Bacillus subtilis*, *Staphylococcus aureus*, and *Streptococcus pyogenes*), and Cyanobacteria (*Synechococcus elongatus*) as well as the model archaea *Sulfolobus acidocaldarius*. The reference genomes for those 14 species are complete, contain few mobile elements (Figure 1B), and encode 95 antiphage systems across 53 types (Figure 1C,D), with Gabija and Paris following R–M and CRISPR-Cas as most prevalent types. To be able to explore the transcriptional diversity of defense system within species, we complemented this species-level database by selecting 15 diverse *E. coli* genetic backgrounds (Figure 1E), including 12 natural isolates from the Picard collection^24^ and three model strains (BW25113, BE, and BL21). We resequenced all strains to obtain complete or near complete genomes, from which we identified a number of potential mobile genetic elements (Figure 1F) as well as a total of 141 antiphage systems spanning 52 types (Figure G,H). To explore antiphage defense transcription across clinically and ecologically relevant settings (see Methods), we sequenced the transcriptomes of *E. coli* strains across six environmental conditions (Lennox LB 37°C, Lennox LB 20°C, Lennox LB + 10g/L NaCl, Lennox LB with pH=8.75, Lennox LB with pH=5.25, and M9 Minimal medium + 2% glucose) as well as two physiological states (exponential phase and overnight cultures as a proxy for stationary phase, see Methods) (Figure 1I,J). We aim to leverage this data to explore transcription patterns of defense systems as a proxy for their expression levels, whose variation across conditions may shed light on their regulation.

**Figure 1:**
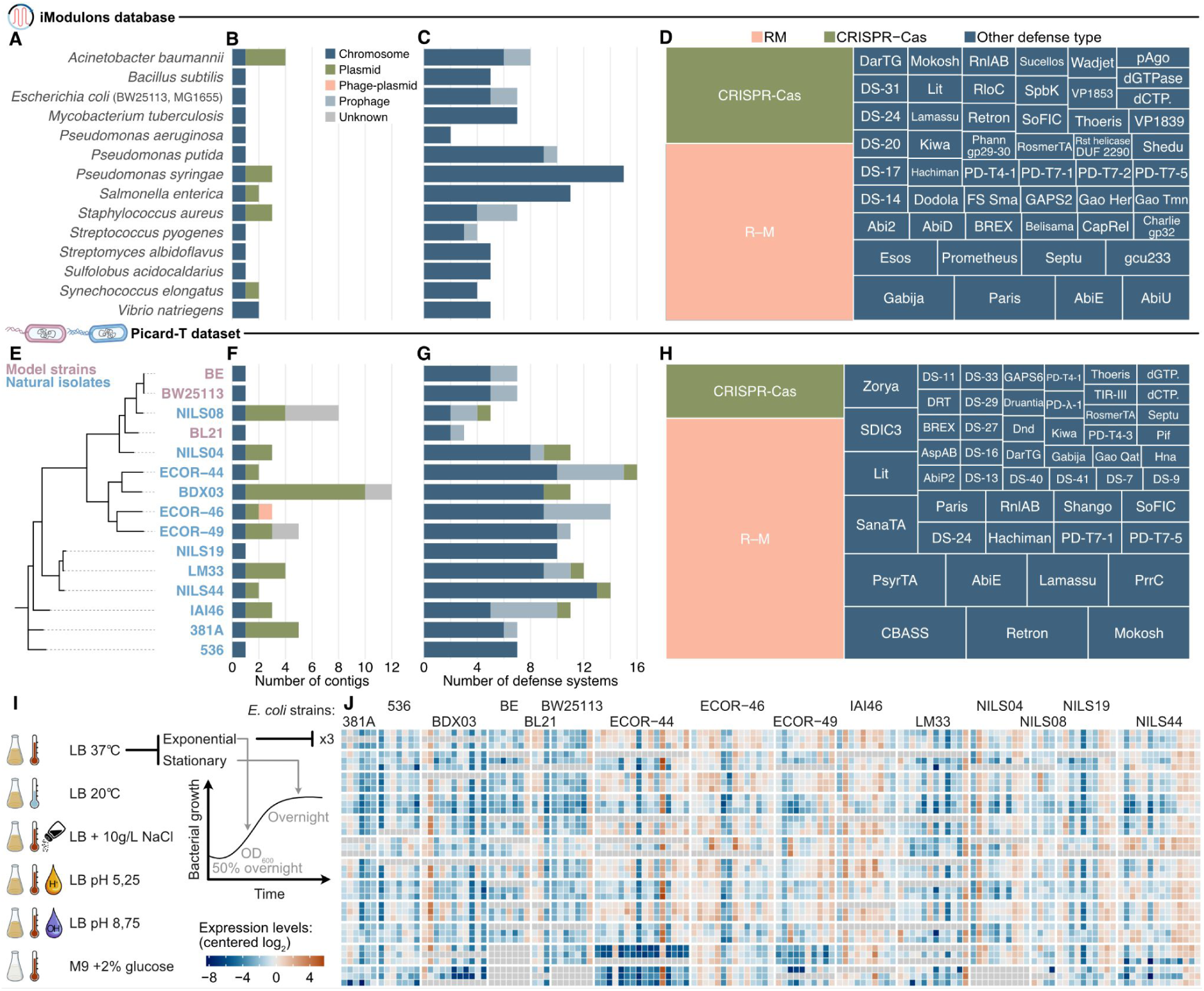
A multi-scale transcriptomic framework to study antiphage system expression. A: Overview of the model species included from the iModulons database and included in this work. B: Number of contigs per reference genome, including MGE predictions (iModulons database). C: Number of the defense systems per reference genomes and their genomic location (iModulons database). D: Overall distribution of the defense systems across types in the iModulons database. E: Phylogeny of the 15 *Escherichia coli* strains selected for this study for the Picard-T dataset. F: Number of contigs per newly resequenced genome, including MGE predictions (Picard-T dataset). **G:** Number of defense systems per strain and their genomic location (Picard-T dataset). H: Overall distribution of the defense systems across types covered by the *E. coli* strain collection of the Picard-T dataset. I: Description of the experimental setup to generate the Picard-T dataset. The 15 isolates were tested in six different conditions, at two different growth stages, and in three replicates leading to the generation of 540 RNA-seq samples. J: Overview of the expression data across the 141 defense systems across the newly generated RNA-seq dataset after quality control of the sequencing data (Methods).

### Antiphage genes are expressed at moderate but variable levels

We sought to leverage this combined transcriptomic data to capture native expression levels of antiphage systems across a range of conditions and hosts. To enable those comparisons we used both quantitative (Centered-Log_2_ Ratios — CLRs) and ordinal (ranked in percentile) transcription levels (Figure 2A, Suppl. Figure 1, Methods). To illustrate the CLR scale and its relationship to ranked transcription levels, we first focused on the model strain *E. coli* BW25113 across the Picard-T dataset (Figure 2A). To further contextualize transcription levels, we show that ribosomal proteins (among the most expressed genes) are typically found in the top 5% most expressed genes with CLRs ranging above 4, while the lac operon (repressed genes) is found in the bottom 20% with CLRs typically below-2. In addition, the ftsYEX operon (strongly expressed genes involved in cellular division^25^) is found in the top 25% with CLRs typically around 1.1–1.8. Antiphage system genes fall in between with moderate transcription levels: we found they are typically in the bottom 50% with a median CLR below 0. This pattern is consistent in the 15 *E. coli* strains across the Picard-T dataset, both on quantitative (Figure 2B) and ordinal (Figure 2D) scales, as well as throughout the 14 model species and associated conditions in the iModulons database (Figure 2E). The moderate expression levels of antiphage genes across strains and species could reflect the autoimmune risks or metabolic costs associated with defense genes^7^.

**Figure 2:**
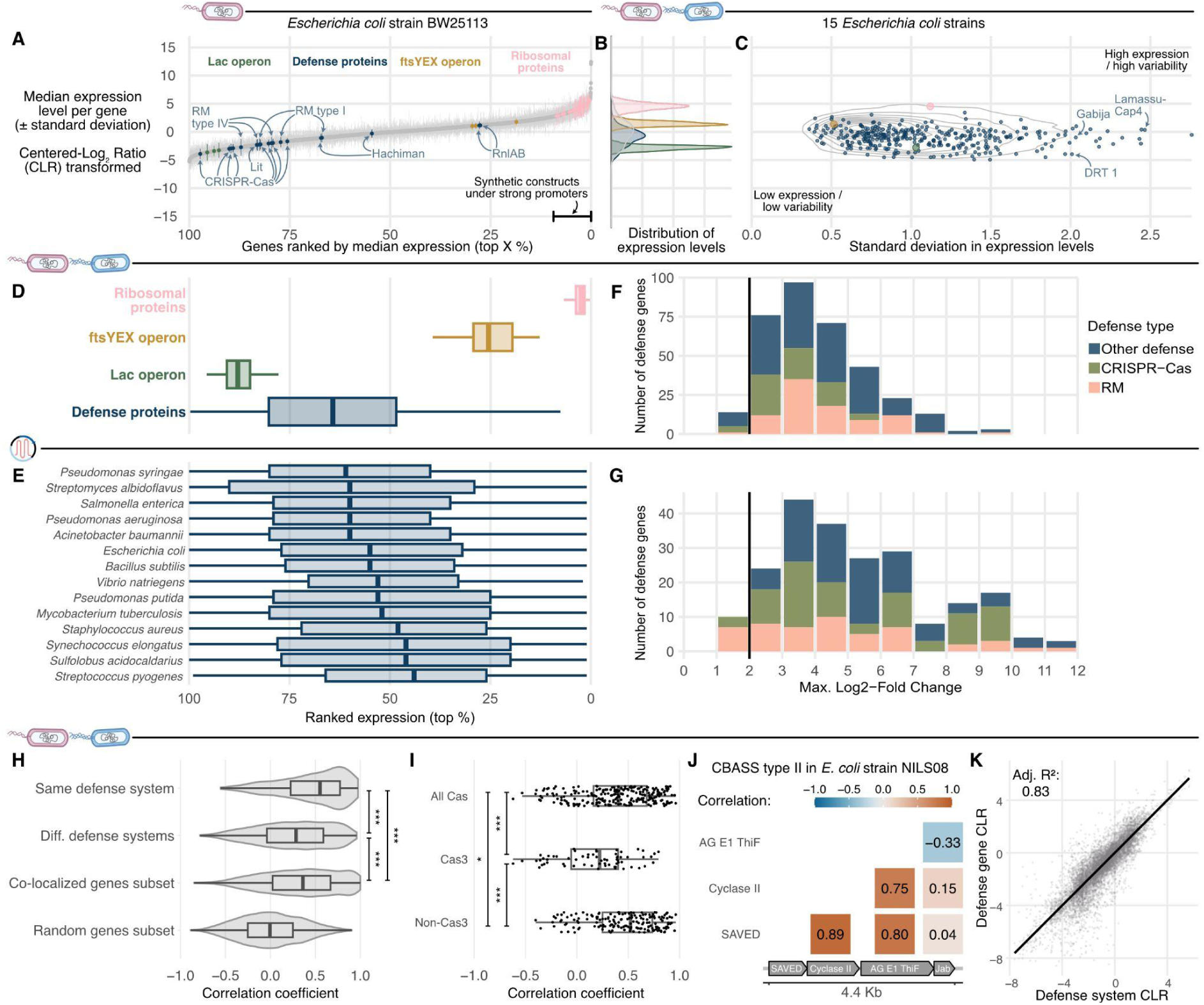
Expression levels and variability of defense systems. A: Relationship between the median expression levels (Centered-Log2 Ratios — CLRs) and ranked expression (in percentile) of *E. coli* BW25113 genes across the newly sequenced transcriptomes (Picard-T dataset). CLRs are centered in 0, which captures the average gene transcription level in a sample, and each unit change from 0 represents a log2-fold change (Log2FC) increase or decrease per unit. All genes are displayed in grey and specific gene sets are highlighted with colors. Synthetic constructs under strong promoters depict the typical range of transcription levels used for molecular characterisation. Similar plots available for all strains and species (Suppl. Figure 2 and 3). B: Distribution of the expression levels of the different gene sets across the Picard-T dataset. C: Relationship between the median expression levels and standard deviation of defense genes in the Picard-T dataset. D: Ranked expression levels of specific genes sets across the Picard-T dataset indicates defense genes are overall expressed at moderate levels. E: Ranked expression levels of defense genes across de iModulons database indicates this moderate expression is conserved across species. F: Maximal log_2_FC observed for each individual defense gene across all samples of the Picard-T dataset. G: Maximal log_2_FC observed for each individual defense gene across the iModulons dataset. H: Correlation of defense genes within and between defense systems. A random subset of genes with colocalizatioπ to capture that of multi-gene defense systems is used as control. A random subset of 4500 genes sampled across the 15 strains is displayed as reference. I: Correlation of CRISPR-Cas genes split by categories: All includes all comparisons, Cas3 includes all comparisons with at least one Cas3 gene, Others include all comparisons without any Cas3 genes. J: Example of correlation between genes of the same defense system. The Jab-domain containing gene of this CBASS type II system is poorly correlated to the other genes. K: Defense gene expression strongly correlates to the median gene expression of the defense system.

While expressed only moderately, we found defense genes to have variable transcription levels across the sampled conditions (Figure B,C,D,E), with strong system-specificity (Figure C). For instance, in the Picard-T dataset, Gabija genes encoded by *E. coli* strain ECOR-49 are expressed at low (average CLR around-1 for both genes) but variable (standard deviation across conditions of 1.6 and 2) levels. Similarly, genes of the Lamassu-Cap4 system encoded by *E. coli* strain NILS44 were expressed on average at CLRs between-0.8 and 0.3 with a standard deviation of ranging from 1.6 to 2.4 across conditions. By contrast, the CRISPR-Cas systems encoded by model strains (BW25113, BE) were found to be both lowly expressed (average CLRs from-4 to-1.5) and with little variation (standard deviation below 1 across conditions). To further quantify this variability, we explored the maximum fold-change of each defense gene across the Picard-T dataset and the iModulons database.

In the Picard-T dataset, all defense genes had a maximum fold change of at least two (Log_2_FC≥1) and 96% of them of at least four (Log_2_FC≥2)—with the restriction enzyme of a type II R–M system of *E. coli* strain ECOR-44 displaying a >800 fold change between two conditions (Figure 2F). We found similar patterns across the iModulons database, with all defense genes displaying at least a two fold change (Log_2_FC≥1) in expression (Figure 2G). In addition, the maximum fold change distribution was not significantly different between defense genes and all genes across both the Picard-T dataset and the iModulons database the (p-values > 0.05, Wilcoxon test, two-side, FDR-corrected), suggesting defense genes are regulated to the same extent as other genes independently of exogenous infection. Together, these results indicate that antiphage systems are typically expressed at moderate but variable levels, including across the conditions covered in both datasets.

### Antiphage systems are complex but cohesive transcriptional units

Having established the expression levels and variability in antiphage genes, we aimed to characterize co-expression patterns within defense systems. In the Picard-T dataset, we found the expression of genes within a defense system were strongly correlated (median 0.58) (Figure 2H). Despite this general trend, we found nuances between defense types or within systems. For instance, CRISPR–Cas systems tend to have lower internal correlation (0.41) compared to the average. However, we found this difference was largely driven by Cas3 (the helicase/nuclease effector of the Cascade complex), which is significantly less correlated (0.22) than the rest of the genes (0.52) (Figure 2I). While these observations may reflect the larger nature of CRISPR-Cas systems^26^ (or the potentially non-functional nature of native *E. coli* CRISPR-Cas systems^27,28^), they also indicate that the effector of a defense system can be under specific transcriptional regulation, potentially to limit self-toxicity. Similarly, we also identified strong correlations between most genes (0.75–0.89) of a CBASS type II operon except one (-0.33–0.15) (Figure 2J). Here, the sensor-transductor-effector complex is highly correlated, but the regulator gene (Jab-domain containing) displays an independent transcriptomic pattern. The Jab-domain protein has been shown to regulate the activity of CBASS systems by (de-)conjugating the CD-NTase^29,30^. We therefore hypothesize that independent transcriptional regulation of this regulator can further modulate the defense activity of the system. Despite such cases, the median expression of multi-gene defense systems still captures very well that of individual genes (R2=0.83) (Figure 2K), independently of the number of genes and similarly to known operons (Suppl. Figure 4). In addition, in the iModulons database, genes from the same system were more correlated to each other than genes from the same iModulon (that is a set of co-expressed genes defined by the iModulons database) (Suppl. Figure 5). We therefore use the median gene expression per defense system for all subsequent defense system-level analyses.

### Environmental, physiological, and spatial gradients drive antiphage systems expression

We next aimed to explore the drivers underpinning variability of antiphage systems expression. To that end, we first leverage the Picard-T dataset to analyze differential expression patterns across both environmental (temperature, salinity, pH, and carbon source) and physiological (growth stages) factors (see Methods). For instance, we found a type II R–M system in the *E. coli* strain NILS44 was upregulated (log_2_FC=4.12) in Lennox LB with increased salinity compared to standard Lennox LB at 37°C. Each individual gene of the operon (Figure 3A) is significantly overexpressed (Figure 3B) although the magnitude of the change varies across the system. We also found examples of defense systems differentially regulated across all conditions covered in our dataset (salinity, temperature, pH, minimal medium, and growth stage). For instance, the Hna defense system encoded by *E. coli* strain NILS08 was upregulated (log_2_FC=1.70) at lower temperatures (Lennox LB 20°C compared to Lennox LB at 37°C, figure 3C). Similarly, pH variation can impact the expression of a PD-T4-3 system encoded by *E. coli* strain IAI46, which was upregulated (log_2_FC=2.06) in Lennox LB at pH=5.25 (compared to standard Lennox LB at 37°C, figure 3E). By contrast the Zorya type I of *E. coli* strain ECOR-44 was downregulated (log_2_FC=-2.22) in minimal medium (compared to Lennox LB at 37°C, figure 3D). Furthermore, we found growth stages can also impact expression with the Thoeris type II of *E. coli* strain 536, which is upregulated (log_2_FC=3.01) in stationary compared to exponential phase (Lennox LB + 10g/L NaCl at 37°C, figure 3F).

**Figure 3:**
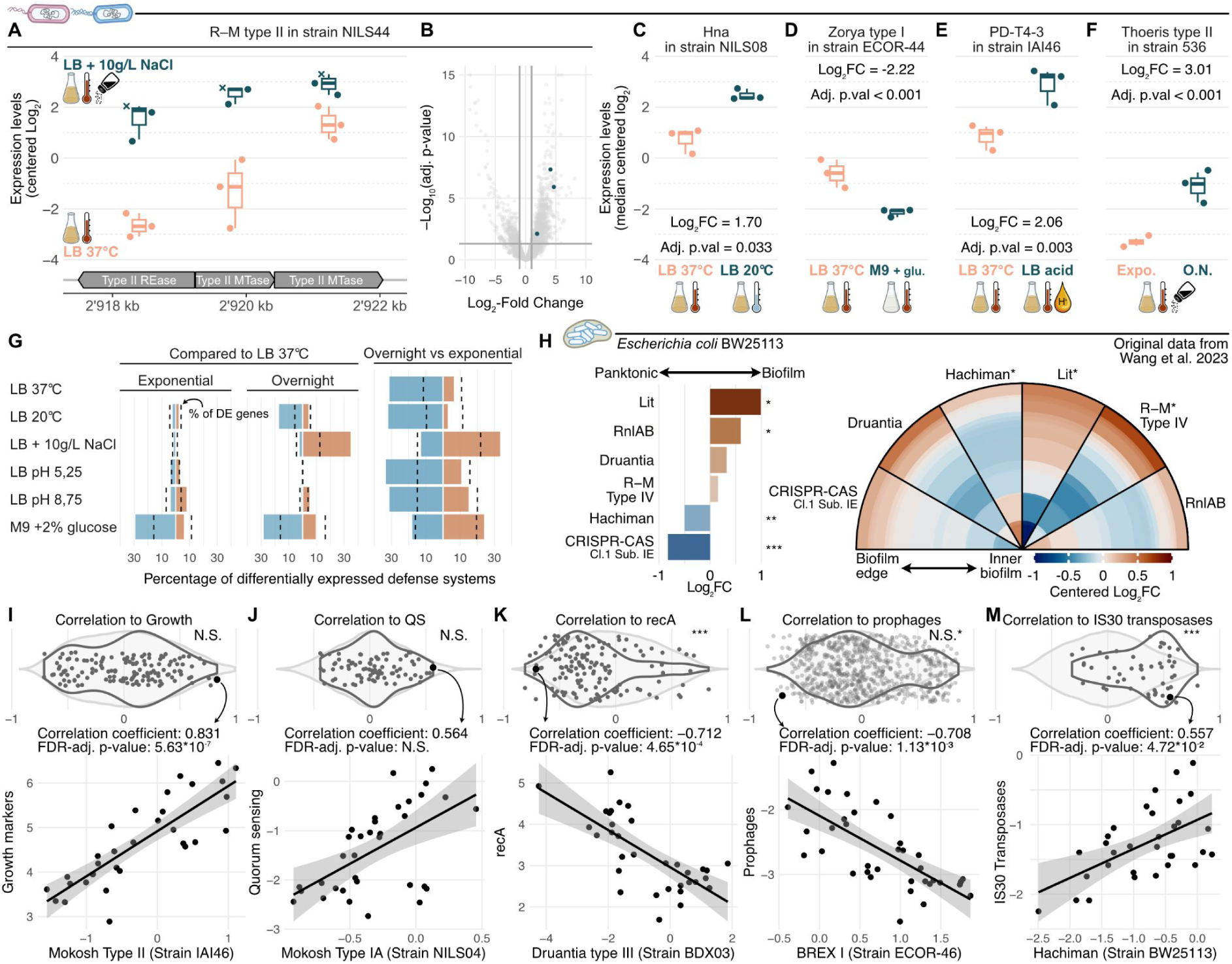
Environmental, physiological, and spatial factors can drive defense transcription. **A:** The individual genes of a R-M type II defense system found in *E. coli* strain NILS44 are overexpressed in LB with NaCI compared to standard growth conditions. Expression levels are shown in CLRs. The sample displayed with crosses is a sample that did not pass QC (see Methods). It is shown here as an indication but excluded from other analyses and statistical tests. B: Each R-M gene shown in A is significantly differentially expressed between the two conditions. Volcano plot based on DESeq2 analyses (Methods). C-F: Median expression levels of defense systems that are differentially expressed across the conditions tested in the dataset. Expression levels are displayed in CLRs, log2FC and significance are based on DESeq2. G: Comparing the number of differentially up-or down-regulated defense systems to the average number of genes reveals key conditions driving differential defense expression. A breakdown across strains indicates these patterns are largely conserved (Suppl. Figure 6) H: Median log_2_FC between planktonic and biofilm samples andsSpatial expression patterns of defense systems across the biofilm. Data is displayed as log_2_FC centered individually for each defense system. Asterisks depict significance as defined in the original work^36^. I-M: Expression correlation between defense systems and specific sets of genes (dark grey) related to growth, quorum sensing (QS), DNA damage (recA), prophages, and transposases. The light grey distribution compares the correlation between all genes and the target set of genes. The correlations captured by points highlighted in black are displayed below.

To determine to what extent these observations hold more broadly, we systematically investigated the proportion of up-and down-regulated defense systems across the different conditions of the Picard-T dataset compared to Lennox LB 37°C. For context, we compared it to the overall proportion of the transcriptome that was up-or down-regulated for each condition. The proportion of differentially expressed systems was largely on par with the overall proportion of differentially expressed genes for many tested conditions (*n*=7), showing that differential expression of defense systems in these conditions may reflect an overall transcriptome shift rather than targeted regulation. However, we found a number of conditions leading to notable enrichments in downregulated defense systems, suggesting a defense-specific transcriptional response to those environmental and physiological changes (Figure 3G, Suppl. Figure 6). In particular, minimal medium, lower temperature, or overnight growth compared to the exponential stage lead to 20–30% of the defense systems being downregulated. Growth in minimal medium, at lower temperature, or in stationary phase was associated with reduced transcription of defense systems, highlighting the tight coupling between metabolic state and defense regulation^31,32^. By contrast, Lennox LB with increased salinity led to a higher transcription of defense systems (>30%), both when comparing overnight Lennox LB + 10g/L NaCl to overnight standard Lennox LB and when comparing overnight to exponential in Lennox LB + 10g/L NaCl. This upregulation may reflect a global transcriptional response to osmotic stress, potentially coupled to an anticipatory response to an increased risk of phage–bacterium interactions in high-salinity environments, where phage adsorption to bacterial surfaces is known to be enhanced^33,34^, by screening electrostatic repulsion^35^.

Since discrete variations in environmental parameters or physiological state impact defense system expression in all strains of the Picard-T dataset, we aimed to explore the consequences of complex, continuous gradients. Specifically, we set out to test whether spatial gradients could lead to differential expression of defense systems. To that end, we reanalyzed planktonic and spatially resolved biofilm RNA-seq data of *E. coli* BW25113 generated by Wang and colleagues^36^ (Suppl. Figure 7). First, we found that four out of the six defense systems were differentially expressed between planktonic and biofilm states (both Lit and RnlAB were overexpressed while Hachiman and CRISPR-Cas were downregulated, data pooled across time points in the planktonic state and across space in the biofilm state). Second, Lit, R–M, and Hachiman displayed spatial expression patterns across the biofilm: Lit and R–M were more highly expressed at the biofilm edge compared to the inner biofilm, while Hachiman was expressed most highly in the inner biofilm (Figure 3H). Together, these results suggest that antiphage defenses can be regulated along environmental, physiological, and spatial gradients, and can lead to a complex population-level structuring of immunity.

### Defense systems expression correlates with markers of cellular physiology and mobile element activity

While discrete environmental, physiological, or spatial gradients can impact defense system expression, whether this is a direct regulation or an indirect response to associated metabolic changes remains open. We posit that co-expression between defense systems and target sets of marker genes across the Picard-T dataset may reflect such complex relationships. For instance, we so far only explored growth as a discrete variable based on two time points (exponential and overnight), yet these two time points may capture very different physiological and transcriptomic states across minimal medium, standard Lennox LB, or different temperatures. We therefore leveraged marker genes that positively correlate with bacterial growth^37^ to map the relationship between defense system expression and growth state continuum (Figure 3I). While the correlations between defense systems and growth markers were not significantly different than those of all genes, we identified specific systems that were strongly correlated to growth markers, both positively, such as a Mokosh type II (encoded by *E. coli* strain IAI46, Figure 3I), or negatively, such as a DRT type 1 system (encoded by *E. coli* strain NILS44, data not shown). Since previous work has shown that defense systems can be regulated by quorum sensing (QS) ^14,19,20^, regulated by stress response^38^, or activated by DNA damage^22,39^, we applied a similar approach to explore the correlation between defense systems and the lsr operon (Figure 3J) as well as SOS response markers (Figure 3K), respectively. We did not find any significant association between QS and defense systems, both across all or individual defense systems (Figure 3J). Although we cannot rule out that additional conditions would capture QS regulation or that specific, unrepresented systems may be regulated by QS, the Picard-T dataset does capture substantial variation in QS marker transcription (log2FC≥2) and suggests that QS-based regulation of antiphage systems is not a systematic feature. By contrast, we found the expression of defense systems was systematically negatively correlated to recA transcription levels (Figure 3K), suggesting that antiphage defenses are repressed during the SOS response.

We further hypothesize that induction of prophages or other mobile-genetic elements between the conditions of the Picard-T dataset^40^ may drive the observed variation in defense systems transcription. To test this hypothesis, we first assumed that the expression levels of prophage structural proteins (e.g., head, tail, capsid) are markers of prophage induction. While most prophage markers were expressed at low levels (median CLR of-1.4) consistent with repression in tested conditions, we did identify prophages with systematically high expression levels (median CLRs > 1): the highest, at median CLR of 2.2 being the phage detected in the *E. coli* strain ECOR-46 assembly, or with sporadically high expression in specific samples, for instance in *E. coli* strain NILS44 with a maximum CLR of 6.5 (Suppl. Figure 8). Even though most prophages are generally repressed, every strain has at least one sample where one of its prophages is upregulated, with expression at least 4-fold (log₂FC ≥ 2) above that prophage’s usual baseline. Together, these results suggest that, even within this limited range of conditions sampled here, bacterial hosts are likely to face prophage inductions. However, correlations between defense systems expression and individual prophages were not significantly different compared to all other genes, despite a clear set of strong positive correlations (Figure 3L) driven by *E. coli* strain ECOR-44. While the *E. coli* strain ECOR-44 encodes many prophages (*n*=10), they do not appear strongly induced, or only rarely. The strong correlations observed are not driven by co-localization (most of the defense systems correlate to prophages they are not part of) but rather driven by a general metabolic shift between minimal medium and other conditions. However, some defense systems display the opposite pattern. For instance, the chromosomal BREX system of *E. coli* strain ECOR-46 is negatively correlated to a prophage located millions of base pairs away (Figure 3L). Similarly, by exploring the correlation between defense systems and plasmids conjugation machinery, we did not find a systematic pattern but specific positive or negative correlations (Suppl. Figure 8). By contrast, we found defense systems were significantly more positively correlated to IS*30* transposase transcription compared to all other genes (Figure 3M). Together, these results show that transcription levels of specific defense systems can correlate with MGE activity.

### Genetic organisation shapes defense system expression

To unwind the genetic basis of defense system regulation, we then set out to explore expression patterns of similar defense systems across strains (Picard-T dataset) at three different levels: systems with high nucleotide identity (>90%), systems with the same defense subtypes, and systems with the same defense types. While the expression of defense systems from the same type (but different subtypes) was not significantly different from systems from different types, the correlation was significantly higher within subtypes and even more so between systems with high nucleotide identity (Figure 4A). This suggests that defense system regulation can be conserved between strains, although some highly similar systems appear to have uncorrelated expression. In addition, we found correlation patterns can be highly variable between defense system subtypes: for instance PsyrTA as well as Mokosh type II defense systems tend to be expressed similarly between strains, while Retron IC or CBASS type I defense systems were poorly correlated between strains (Figure 4B). For instance, the PsyrTA defense systems of strains *E. coli* strain ECOR-46 and strain NILS44 displayed strongly correlated expression patterns (correlation coefficient of 0.87) (Figure 4C). By contrast, the CBASS type I defense systems of *E. coli* strain NILS04 and strain NILS44 had uncorrelated expression patterns despite being >99% identical at the nucleotide level (Figure 4D). Interestingly, the upstream regions are relatively well conserved in both cases (97% and 96% nucleotide identity across 100 bp for PsyrTA and CBASS, respectively). However, the two PsyrTA systems are found in similar genomic context while the two CBASS systems are found on different plasmids with variable gene content. Overall, we found that closely related systems tend to display conserved expression patterns between genetic backgrounds although some nearly identical defense systems still exhibited uncorrelated expression patterns between strains. These observations suggest that regulation of defense system expression can evolve rapidly, providing an additional dimension for evolution to tinker with and generate immune diversity.

**Figure 4:**
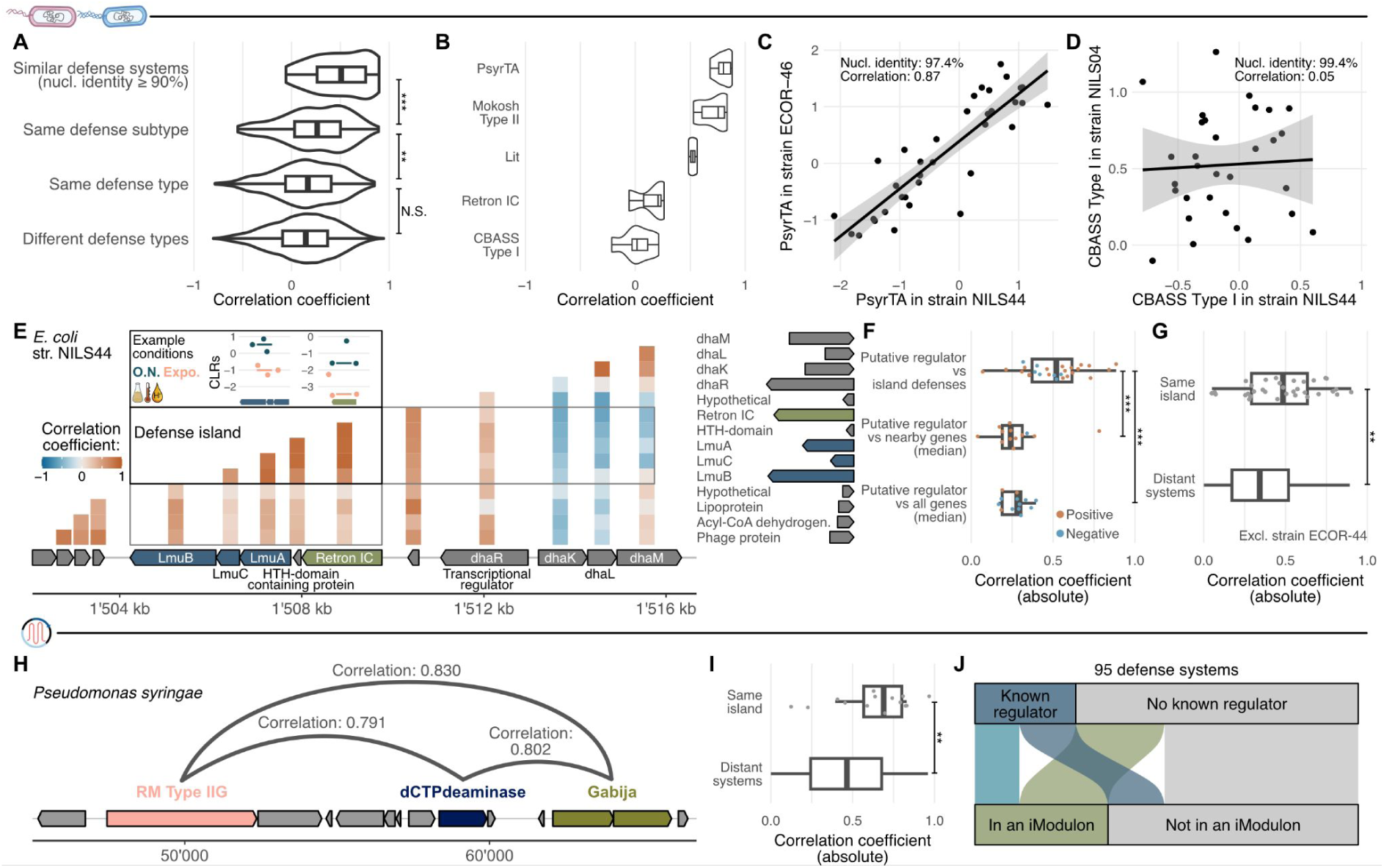
Genetic organisation of defense system regulation. **A:** Expression correlation between defense systems across strains from the Picard-T dataset. The more similar two systems are, the more correlated is their expression. B: Similar expression patterns between related defense systems across strains is system-subtype dependent. **C-D:** Example of two highly similar defense systems with highly and poorly correlated expression patterns across strains, respectively. **E:** Example of a candidate defense island in *E. coli* strain NILS44 with leading and trailing genes displayed along with the expression correlation between genes. The inset displays the median expression levels (CLRs) of the two defense systems across two example conditions to illustrate the correlation. F: Across all 32 candidates defense highlands, 25 had putative regulators. Those regulators are significantly more correlated to the defense systems from the island than to other genes. G: Overall, defense systems from the same island are more correlated than distant defense systems. Comparison done excluding *E. coli* strain ECOR-44 (see Methods). H: Example of co-localized defense systems in Pseudomonas syringae and their pairwise correlation coefficients. I: Correlation coefficients between pairs of co-localizing are higher than between genomically distant defense systems. **J:** Almost half of the 95 defense 95 defense systems in the iModulons database have at least a gene with a known regulator or that belongs to an iModulon.

Defense systems are often found to co-localize on so-called defense islands^41^, which is suggested to lead to co-expression due to local regulators^42–46^. Since genes from different defense systems have higher correlations (median 0.35) than random genes (Figure 2H), we hypothesize this may be a result of island-level regulation and/or shared global regulators^47^.

To test the first hypothesis, we explored the correlation patterns of defense systems belonging to the same defense island (Picard-T dataset, Methods). We started by focusing on a defense island encoded by the *E. coli* strain NILS44. The island is composed of a Lamassu and a Retron IC defense systems separated only by an unknown, HTH-domain containing protein (Figure 4E). We found the expression of the five genes to be highly correlated with each other, by contrast with other neighbouring genes (Figure 4E). As HTH-domains are often involved in transcriptional regulation^48^ and local regulators tend to be transferred with their target genes^49^, we posit that the unknown protein may be involved in the co-regulation of the defense island. By inspecting the 32 defense islands identified across 14 *E. coli* strains, we identified putative regulators in 13 of them (Methods). The expression of those regulators was significantly more correlated to defense systems belonging to the island compared to nearby, non-defense genes or all other genes (Figure 4F, Suppl. Table 1). Finally, we found defense systems within the same island showed stronger correlations in their expression patterns compared to distant defense systems (Figure 4G, Methods) — although pairs of distant defense systems can still be highly correlated. We then explored defense islands across species by focusing on the iModulons database. We found *Pseudomonas syringae* to harbour a defense island that includes three defense systems: a type IIG R–M system, a dCTPdeaminase, and a Gabija defense system (Figure 4H). The expression patterns of the three defense systems were highly correlated (0.79–0.83). We subsequently generalized our investigation to the nine defense islands identified across the 14 species (leading to 13 pairs of defense systems). While the data is sparse, defense systems on the same island were significantly (p-value=0.002, wilcoxon-test) more co-expressed (0.70) compared to non-colocalizing systems (0.43) across the iModulons database (Figure 4I), supporting conclusions based on the Picard-T dataset. In addition, we found that nearly half of the defense systems in the iModulons database (n=95) had at least a gene with a known regulator and/or member or an iModulon^50^ (group of co-expressed genes). Local, island-level regulation is further supported by defense genes associated with iModulons of genomic islands (for instance in *Staphylococcus aureus*)^51^. By contrast, we also found defense genes associated with global regulators in transcriptional regulatory networks, including regulators of carbohydrate metabolism (Ccpa, CRP)^52,53^ as well as the SOS response (LexA)^54^, both supporting previous reports^21,39^ as well as our conclusions based on the Picard-T dataset. Together, these results suggest that co-localization may indeed be associated with co-regulation and that both island-level and global regulators are likely to shape the expression of the antiphage repertoire.

### Observed variation in transcription levels has functional relevance

Having established that the expression of defense systems can vary across environmental conditions, growth stages, and genetic backgrounds, we hypothesize that this information can be leveraged to characterize defense systems at native expression levels. For instance, many *E. coli* strains encode a CRISPR-Cas class 1 defense system^55^. Yet, these are largely considered to be non-functional in laboratory conditions^27,28,56^. While the transcriptional repressors of this system have been identified enabling engineered functional CRISPR-Cas system in model strains ^14^, to the best of our knowledge a functional native CRISPR-Cas system has not yet been validated in *E. coli* natural isolates in lab conditions. We therefore explored the expression of CRISPR-Cas Class 1 Subtype I-E in our Picard-T dataset, aiming to find strain and growth conditions that may provide evidence of activity. First, we found that the expression was highly variable across strains. While most of them expressed the CRISPR-Cas system at very low levels, ECOR-44 and even more so ECOR-46 expressed their systems at higher levels overall (Figure 5A). In addition, we found that the CRISPR-Cas defense system in ECOR-46 was more expressed in lennox LB at 20°C compared to other conditions (Suppl. Figure 9). Therefore, by switching from the model strain *E. coli* BW25113 in standard growth conditions (Lennox LB 37°C) to a natural strain in optimised growth conditions, we can increase the expression of CRISPR-Cas by almost ten folds (up to CLR=0.78) (Figure 5B). We therefore tested the ability of the CRISPR-Cas systems to limit transformation rates of a plasmid bearing a genomically encoded CRISPR spacer for both strains and across both conditions (Suppl. Figure 10). We found that only the CRISPR-Cas system of the *E. coli* strain ECOR-46 in optimised conditions was able to significantly reduce the transformation rate by more than seven fold. Beyond validating a functional CRISPR-Cas system in *E. coli* under lab conditions, these results demonstrate how unwinding the transcriptional regulation of defense systems across conditions and genetic backgrounds can help guide the characterization of defense systems at physiologically relevant expression levels.

**Figure 5:**
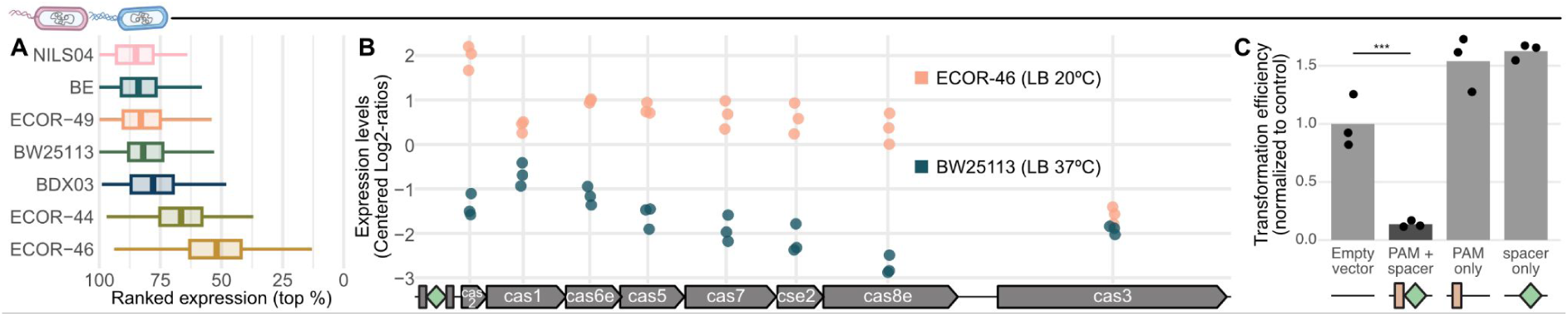
Functional relevance of differential expression in native systems. **A:** Median rank-normalized expression (percentile) of CRISPR-Cas Class 1 Subtype IE in the Picard-T dataset the variability in expression levels, with E. coli strain ECOR-46 expressing CRISPR-Cas the highest although still at moderate levels. B: Per-gene expression levels of the CRISPR-Cas locus in model conditions *(E. coli* BW25113 in LB 37°C) and optimised strain and conditions *(E. coli* ECOR-46 in LB 20°C). **C:** In optimised conditions *(E. coli* ECOR-46 in LB 2Ũ°C) the transformation efficiency of a spacer-bearing plasmid was 7 fold lower compared to an empty control.

## Conclusion

Together, our work provides a systematic assessment of antiphage systems transcription patterns, showing that they are moderately but variably expressed along environmental, physiological, and spatial gradients. Building on these results, we explored the co-expression between defense systems and markers of cellular physiology as well as mobile genetic element activity, identifying both systematic (SOS response, IS*30* transposases) and specific (growth markers, individual prophages) associations. We also shed light on the genetic organisation of prokaryotic immune regulation, including island-level co-expression with candidate local regulators as well as variability across strains. We further demonstrate the functional relevance of expression levels variation across strains and conditions.

In eukaryotes, the immune response heavily relies on transcriptomic upregulation triggered by the infection. By contrast, our results add to previous work on individual systems and suggest that a plethora of phage-independent factors can regulate antiphage defense systems. This complex regulation may prime bacteria to better resist infections under specific ecological conditions as well as mitigate the fitness cost associated with the antiphage repertoire of prokaryotes. This complex regulation can also lead to a population-level organisation of defense system expression (as seen by the spatial organisation in a biofilm), raising a number of questions on the single-cell heterogeneity in defense system expression levels.

## Code and data availability statement

The code used for the analyses performed in this study is accessible on GitHub (https://github.com/LucasPaoli/antiphage-expression). Intermediary data produced during the analysis is available on Zenodo (https://doi.org/10.5281/zenodo.17929710). The primary data (raw RNA-seq) will be deposited at ENA before publication.

## Acknowledgements

We thank the Bioinformatics and Biostatistics Hub as well as the Biomics Core Facility of the Pasteur Institute for their support. L.P., R.L., A.L., F.T., and A.B. are supported by the European Research Council Starting Grant (PECAN 101040529) and core funding from the Pasteur Institute. L.P. is supported by a European Molecular Biology Organization Postdoctoral Fellowship (EMBO ALTF 100-2023). We thank all the members of the MDM Lab as well as François Rousset for their feedback on the manuscript.

## Methods

### Generation of the Picard-T dataset

#### Strain selection

We selected twelve strains from the Picard collection^24^, representative of the phylogenetic diversity of *Escherichia coli* and of the diversity of known anti-phage defense systems, and included the BW25113, BE, and BL21 model strains.

#### Genome sequencing

For each strain, we re-sequenced the genome using a hybrid platform at an external service provider (Plasmidsaurus) to ensure complete or near-complete references for downstream analysis. Genomes were annotated with bakta^57^ (v1.11.4) to identify and annotate genes and genomic features, genomad^58^ (v1.11.1) to identify and annotate mobile genetic elements, and defensefinder^59^ (DefenseFinder v2.0.1, macsyfinder v2.1.4, and Defensefinder model v2.0.2) to identify antiphage systems.

#### Culture conditions

We cultured the fifteen bacterial strains in six different growth media to exponential and stationary growth phases, independently of any external experimental phage predation or other mobile genetic elements (MGEs).

Our reference medium was Lennox Broth (BD Difco, 240230; osmolarity 240 mOsm/L, pH 7.30, tryptone 10 g/L), incubated at 37 °C. We then varied one parameter at a time from this reference condition: (1) incubation temperature (20 °C); (2) medium osmolarity (adding 10 g/L NaCl, yielding an osmolarity of 580 mOsm/L); (3) acidic pH (adjusted to pH 5.25 by adding HCl); (4) alkaline pH (adjusted to pH 8.75 by adding NaOH); and finally (5) the energy substrate, using minimal M9 salts (BD Difco, 248510) supplemented with 20 g/L glucose as the energy source, with added MgSO₄ and CaCl₂. These abiotic conditions were selected to capture two major ecological niches of *Escherichia coli*: the gut and extra-intestinal infection sites of endothermic animals versus environmental water and soils, primarily differing in temperature, as well as three key physico-chemical parameters that vary at infection sites. Using urine as a particularly complex and physiologically variable example, these parameters include osmolarity (which can range physiologically from 60 to 1200 mOsm/L), pH (which can vary from 4.5 to 8, a range largely overlapping with that permissive for E. coli growth, approximately pH 5 to 9), and the energy source. For the latter, we compared two distinct energetic and metabolic regimes in *E. coli*: a rich, peptide-based energy supply provided by tryptone in Lennox LB medium, and a defined, glucose-based energy source in minimal M9 medium.

Bacterial strains were pre-adapted by overnight culture in the target medium in 96-well plates in a microplate reader (Tecan Infinite 200), allowing continuous monitoring of optical density at 600 nm (OD₆₀₀) for each sample. Bacteria were then placed in exponential growth by transferring 2 µL of the overnight culture into 200 µL of the same medium in a fresh 96-well plate using an Integra MINI96 automated pipetting system under a laminar flow hood. Exponential growth plates were incubated in the same microplate reader as the overnight cultures (Tecan Infinite 200) and harvested when the OD₆₀₀ reached 40–60% of that of the corresponding overnight cultures. Overnight cultures were harvested at the time of inoculation of the exponential growth plates. Three biological replicates were performed for each strain in each condition.

#### RNA-seq

After being harvested, the 200 μL of bacterial culture were transferred into 1.5mL Eppendorf tubes (RNAase free) containing 400 μL of RNAprotect Bacteria Reagent (Qiagen, 76506). After homogenization, the Eppendorf tubes were left for 5 minutes at room temperature, then centrifuged (10 minutes, 5000G, 4°C), then the supernatant was eliminated and the pellet preserved at-80°C. Pellets were sent to Genewiz for RNA extraction and sequencing on an illumina short-read platform with 2×150bp reads and a target of 10M reads per sample.

#### Primary data analysis

First, the raw sequencing reads were quality controlled using BBMap (v39.37) to remove remaining adapter sequences (ktrim=r k=23 mink=11 hdist=1), potential PhiX contamination (k=31 hdist=1), and trim and filter low quality reads (qtrim=rl maq=20 maxns=1 trimq=14). Quality-controlled reads were subsequently mapped to references using BWA^60^ (v0.7.18) (mem-a) and only alignments of at 45bp and a read coverage of 80% with an identity ≥97% were retained. Alignments were subsequently processed using FeatureCounts^61^ (subread v2.0.6) (-O --primary --fraction) to generate feature count tables. The number of reads mapped was computed using samtools (v1.17) (view-F 0×904-c) and used to derive mapping rates by comparing to the total number of quality-controlled reads.

To ensure there was no cross-contamination between samples, we first built a pangenome of the 15 *E. coli* strains using CD-hit (v4.8.1) (cd-hit-est-c 0.95-T 48-M 0-G 0-aS 0.9-g 1-r 0-d 0-b 100). We identified gene clusters containing genes originating from a single strain as strain markers. We mapped quality-controlled reads to the pangenome representatives as described above. Strain-specific markers were used to manually check (1) that the sequenced strain matched the sample labels and (2) for potential cross-contamination between samples.

We subsequently aligned the quality-controlled reads to the reference genome of each strain as described above. When exploring mapping rates, we identified a number of samples with large fractions of unmapped reads. After further inspection, we found samples with low mapping rates to have a significant proportion of reads mapping to the human genome (mapped to GCF_000001405.40, as described above). This pattern was highly dependent on the sequencing plate number at the external sequencing provider. We therefore concluded in library contamination at the external sequencing provider, which can decrease the effective sequencing depth of impacted samples but is unlikely to impact the biological results. We therefore filtered out samples with less than five million reads mapped and proceeded with downstream analysis.

### iModulons database

We leveraged the iModulonDB^23^ (iModulons database) to include standardized, high-quality transcriptomic profiles, iModulons, as well as known transcription regulators for model species. Transcriptomic profiles and annotations were matched using reference genomes as follows: *Acinetobacter baumannii* (Precise 139^62^; GCA_000963815.1), *Bacillus subtilis* (Modulome^63^; GCF_000009045.1), *Escherichia coli* (Modulome^64^; GCF_000005845.2), *Mycobacterium tuberculosis* (Modulome^65^; GCF_000195955.2), *Pseudomonas aeruginosa* (Precise 411^66^; GCA_000006765.1), *Pseudomonas putida* (Precise 321^67^; GCA_000007565.2), *Pseudomonas syringae* (Precise 202^68^; GCF_000007805.1), *Salmonella enterica* (Modulome core^69^; GCF_000006945.2), *Staphylococcus aureus* (Precise 165^51^; GCF_000017085.1), *Streptococcus pyogenes* (Modulome^70^; GCA_001021955.1), *Streptomyces albidoflavus* (Modulome^71^; GCF_000359525.1), *Sulfolobus acidocaldarius* (Modulome^72^; GCF_000012285.1), *Synechococcus elongatus* (Precise 300^73^; GCA_000012525.1), and *Vibrio natriegens* (Precise 148^74^; GCF_001456255.1). All reference genomes were re-annotated with bakta (v1.11.4) to identify and annotate genes and genomic features and genomad (v1.11.1) to identify and annotate mobile genetic elements, and defensefinder (DefenseFinder v2.0.1, macsyfinder v2.1.4, and Defensefinder model v2.0.2) to identify antiphage systems.

*Limosilactobacillus reuteri* was excluded from downstream analysis as the genome (GCF_020785475.1) used in the original study^75^ was too fragmented (71 contigs).

### Spatial RNA-seq in an *E. coli* biofilm

We included publicly available data from an external study based on *E. coli* BW25113^36^ to explore the expression of defense systems across spatial gradients within a biofilm setting (Suppl. figure 7). The published expression profiles were transformed into CLRs and the expression of multi-gene antiphage systems was computed as the median of individual genes. Comparison between planktonic between biofilm conditions was performed by comparing planktonic samples across all time points to biofilm samples across the full spatial gradient (Wilcoxon test, two-sided). Log_2_-fold changes were computed by taking the difference between the median of each condition. For spatial patterns, the CLRs of each defense system were centered around the mean CLR of that system in the biofilm.

### Expression data analysis

#### Data normalization and processing

Counts were variance-stabilized using the VST function of DESeq2^76^, normalized by gene length (variance-stabilized counts per kb), and used to compute ranked expression in percentiles as well as Centered Log_2_ Ratios (CLR). In addition, raw counts were used to compute differential expression between conditions using DESeq2. Ranked expression is used for ordinal representation of expression levels, CLRs are used to display quantitative expression levels, compute maximum log_2_-fold changes, and compute correlations between gene or gene set expression profiles, and DESeq2 results are used to compute the significance of differential expression between conditions.

#### Correlations to marker genes

For each strain, we computed Pearson correlation coefficients between defense systems and target gene sets based on CLRs. As a proxy for growth, we used a set of universal single-copy marker genes characterized in *E. coli*^37^, for quorum sensing (QS) we used the lsr operon, for prophages we used structural protein (head, tail, capsid), and for plasmids we used the conjugative machinery (as predicted by conjscan^77^). Per sample, we compute the median expression of each defense system and each gene set to compute the correlations.

#### Identification of defense islands

Defense islands were identified through a two-step process. First, we identified all stretches of antiphage systems separated by less than 30 kb (threshold based on the distribution of all the distances to the next defense system across genomes). We subsequently manually inspected each of these candidate islands (including all genes less than 10 kb upstream or downstream of the first or last system, respectively) and only retained those with consecutive antiphage systems, or if they are separated by only few genes and/or genes with candidate defense annotations (e.g., toxin/antitoxin, DEAD/DEAH box helicase, restriction nuclease, TIR domain). We additionally identified potential regulators based on gene annotations, including WYL or HTH domain containing proteins as well as other annotated transcriptional regulators.

#### Pairwise identity of antiphage systems

To explore expression across similar and different defense systems, we computed pairwise nucleotide identity between defense systems using vsearch^78^ (v2.30.1) (--allpairs_global --acceptall). We grouped defense systems from the most to the least similar as follows: nucleotide identity ≥90%, same subtype, same type, different type (as defined by defensefinder).

### CRISPR-Cas interference assay

For each assay, both *E. coli* strains (ECOR-46 and BW25113) were inoculated from a fresh plate and grown overnight at either 37°C or 20°C. For assays performed at 20°C, cultures were reinoculated, and cells were made electrocompetent by treatment with 10% glycerol. Then, 25 µL of competent cells were transformed with 20 ng of the plasmids listed in Suppl. Table 2. After 2 hours of recovery at 20°C, 50 µL of cells were plated on carbenicillin-containing plates (100 µg/mL) and incubated at 20°C for at least 24 hours. Colony-forming units (CFUs) were counted to determine the transformation efficiency. For assays performed at 37°C, the same protocol was used to prepare electrocompetent cells. Then, 100 µL of ECOR-46 competent cells and 25 µL of BW25113 competent cells were transformed with 20 ng of the same plasmids listed above. After 30 minutes of recovery at 37°C, 100 µL of ECOR-46 cells and 20 µL of BW25113 cells were plated on carbenicillin-containing plates and incubated overnight at 37°C. CFUs were counted to determine the transformation efficiency. The results were normalized to the control for each strain and condition by dividing the CFU/µg of DNA by the mean of the control (empty plasmid). Statistical significance was computed using pairwise t-tests (two-sided) on the normalized data with FDR correction for each strain and condition individually.

## Supplementary tables

**Supplementary table 1** Curated genomic context of the candidate defense islands identified in the Picard-T dataset and iModulons database.

**Supplementary table 2** Results of the CRISPR-Cas interference assays.

## Supplementary figures

**Supplementary figure 1:**
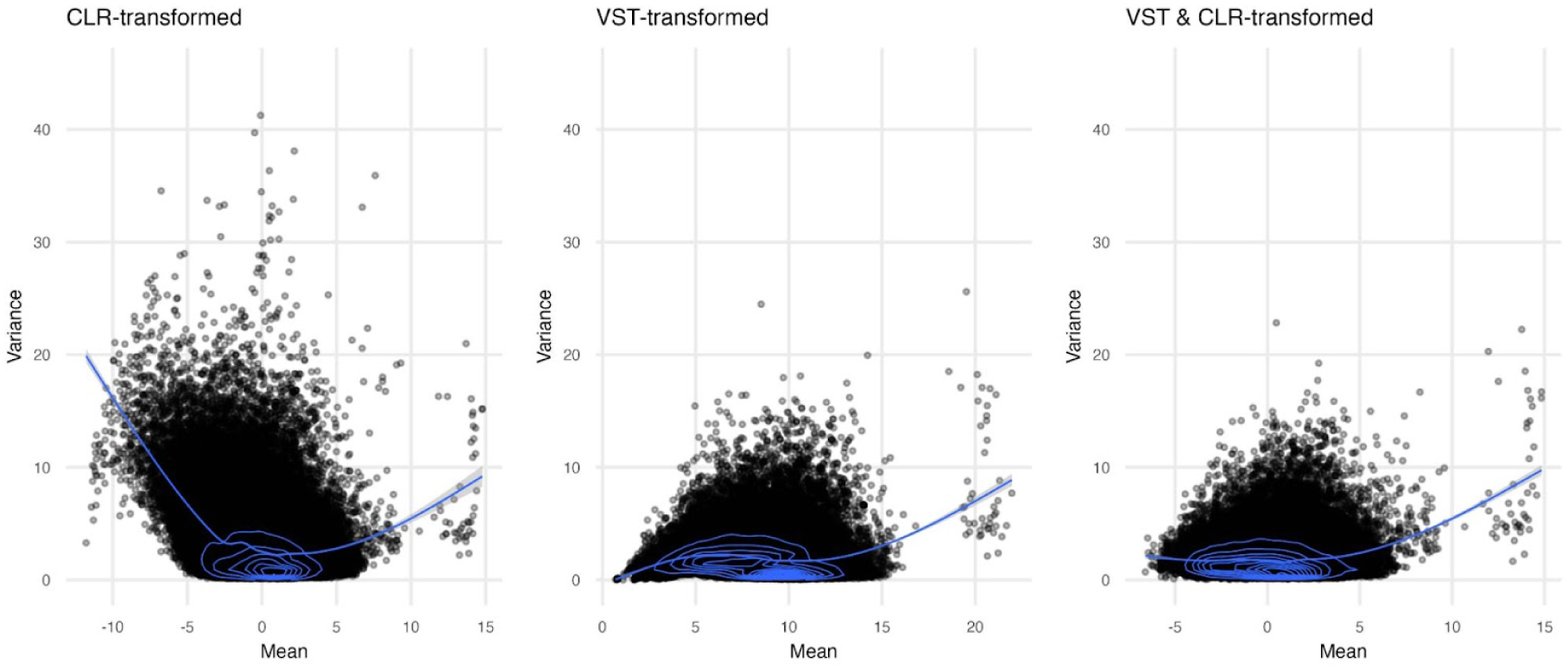
RNA-seq data normalization. Mean–variance relationships across multiple data transformation strategies: centered log-ratio (CLR) transformation alone, Variance Stabilization Transformation (VST) as encoded in DESeq2, and the combination of both. Combining both approaches reduces heteroskedasticity while zero-centering the data, facilitating cross-comparisons and interpretation.

**Supplementary figure 2:**
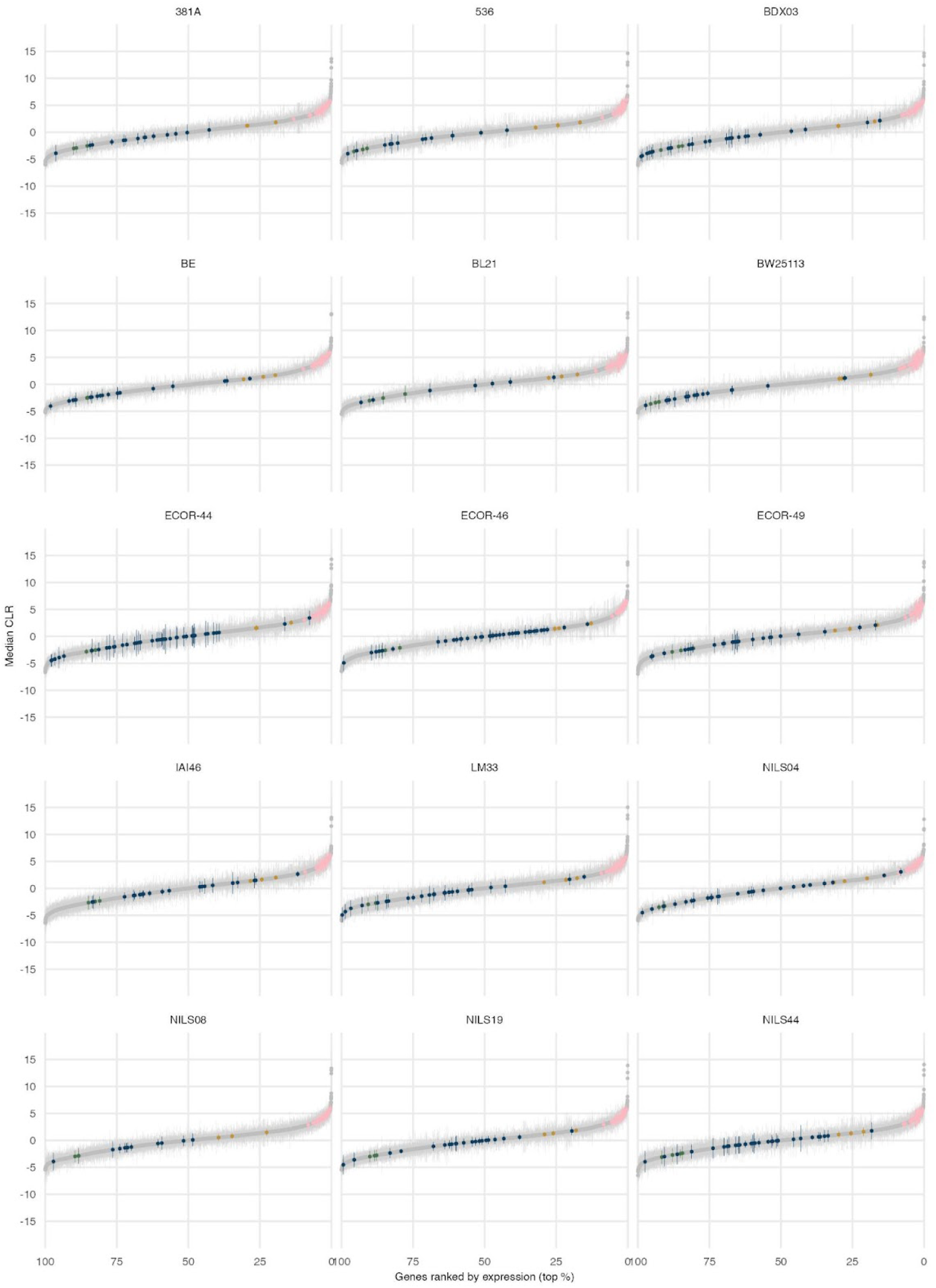
Rank-CLR distribution of genes across E. coli strains. Relationship between the median expression levels (Centered-Log2 Ratios — CLRs) and ranked expression (in percentile) of genes for each strain across the newly sequenced transcriptomes (Picard-T dataset). All genes are displayed in grey and specific gene sets are highlighted with colors as per Figure 2A.

**Supplementary figure 3:**
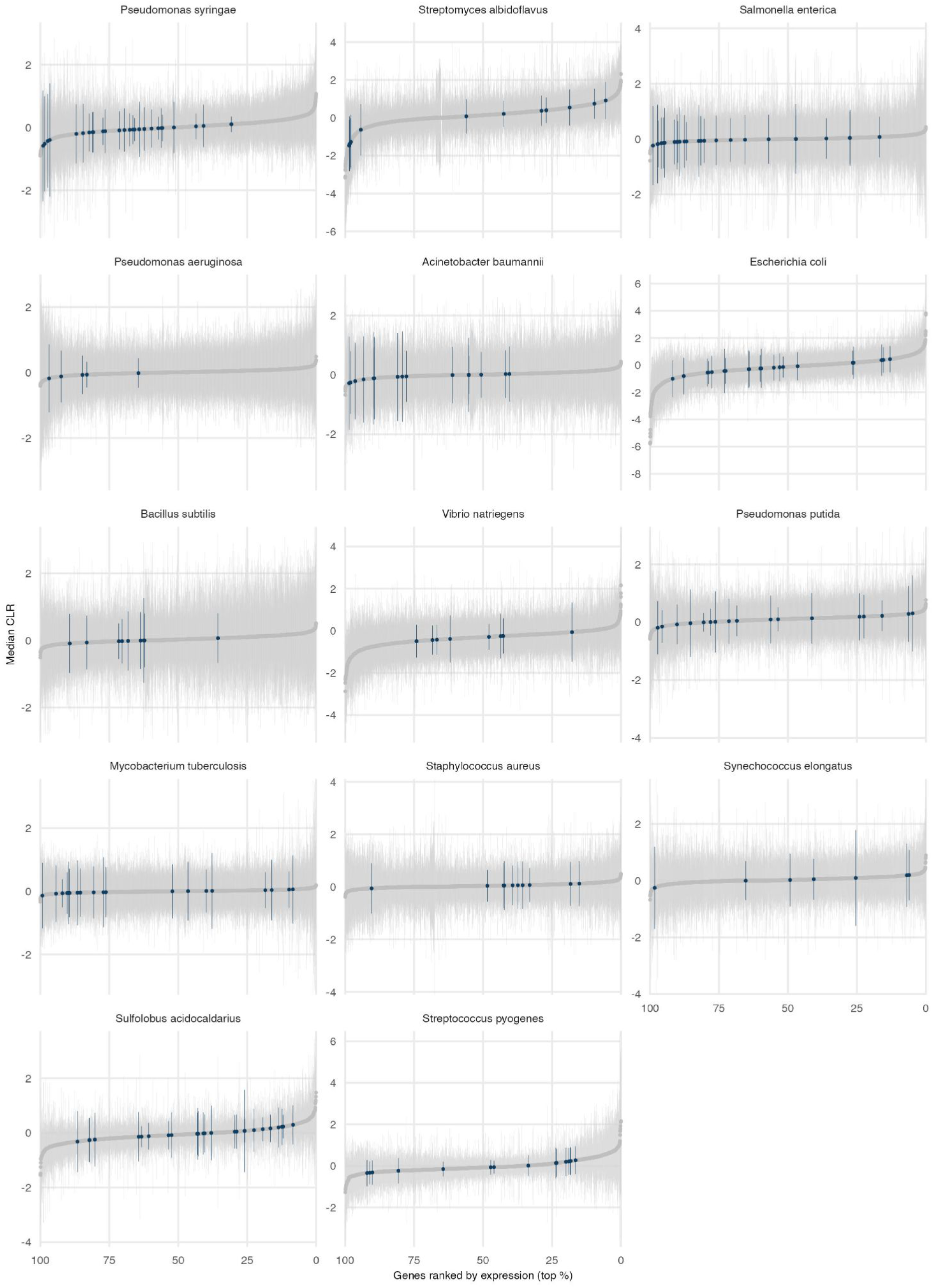
Rank-CLR distribution of genes across iModulons species. Relationship between the median expression levels (Centered-Log2 Ratios — CLRs) and ranked expression (in percentile) of genes for each species in the iModulons database. All genes are displayed in grey and defense genes are highlighted in color.

**Supplementary figure 4:**
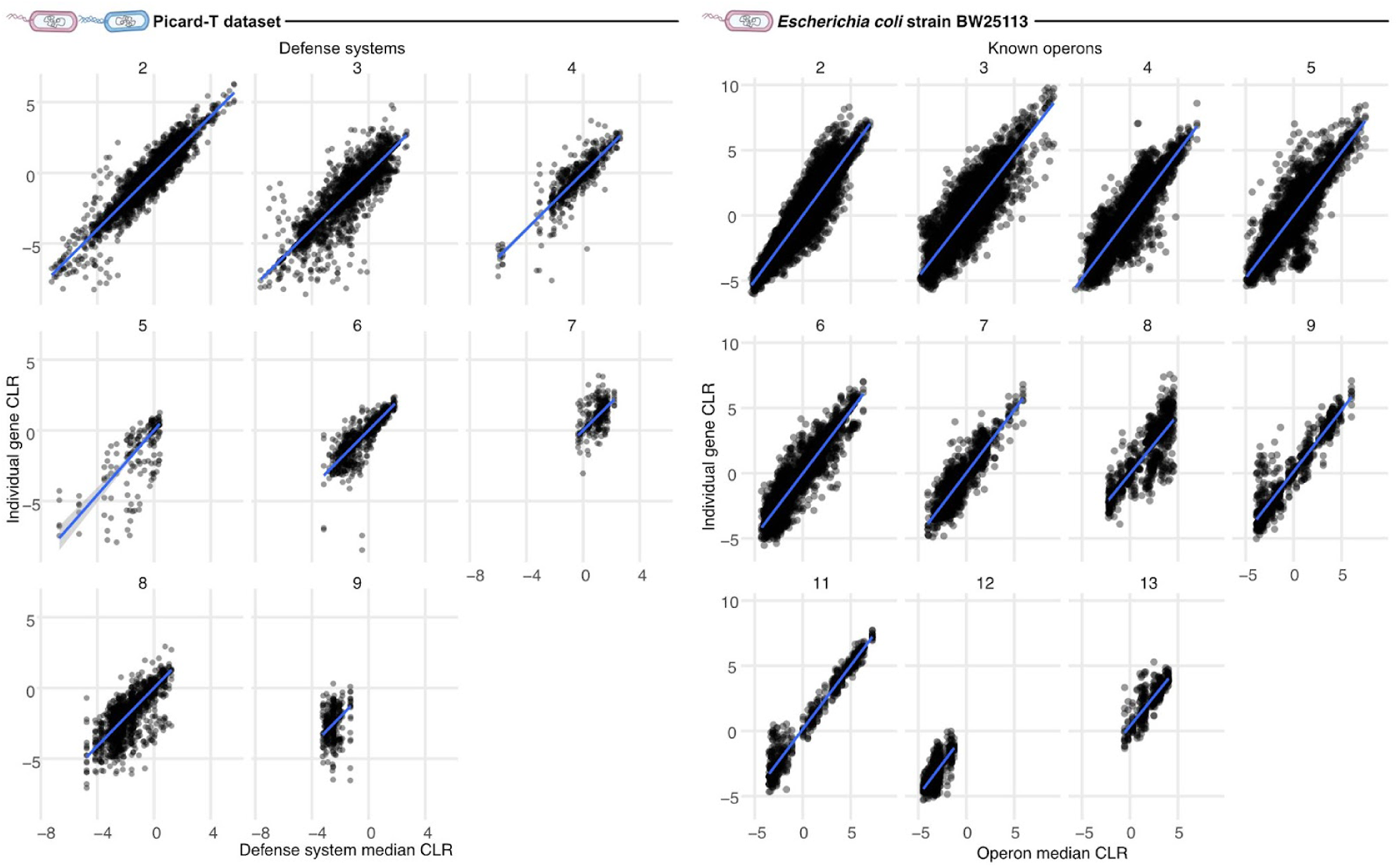
Operon-level correlation of defense systems. The relationships (linear model Y ∼ X + 0) between genes within defense systems (R^2^=0.83) are similar to known operons (R^2^=0.86). The number of genes in the defense system or the operon only marginally impact the linear model, with R^2^≥0.79 across individual defense system or operon sizes.

**Supplementary figure 5:**
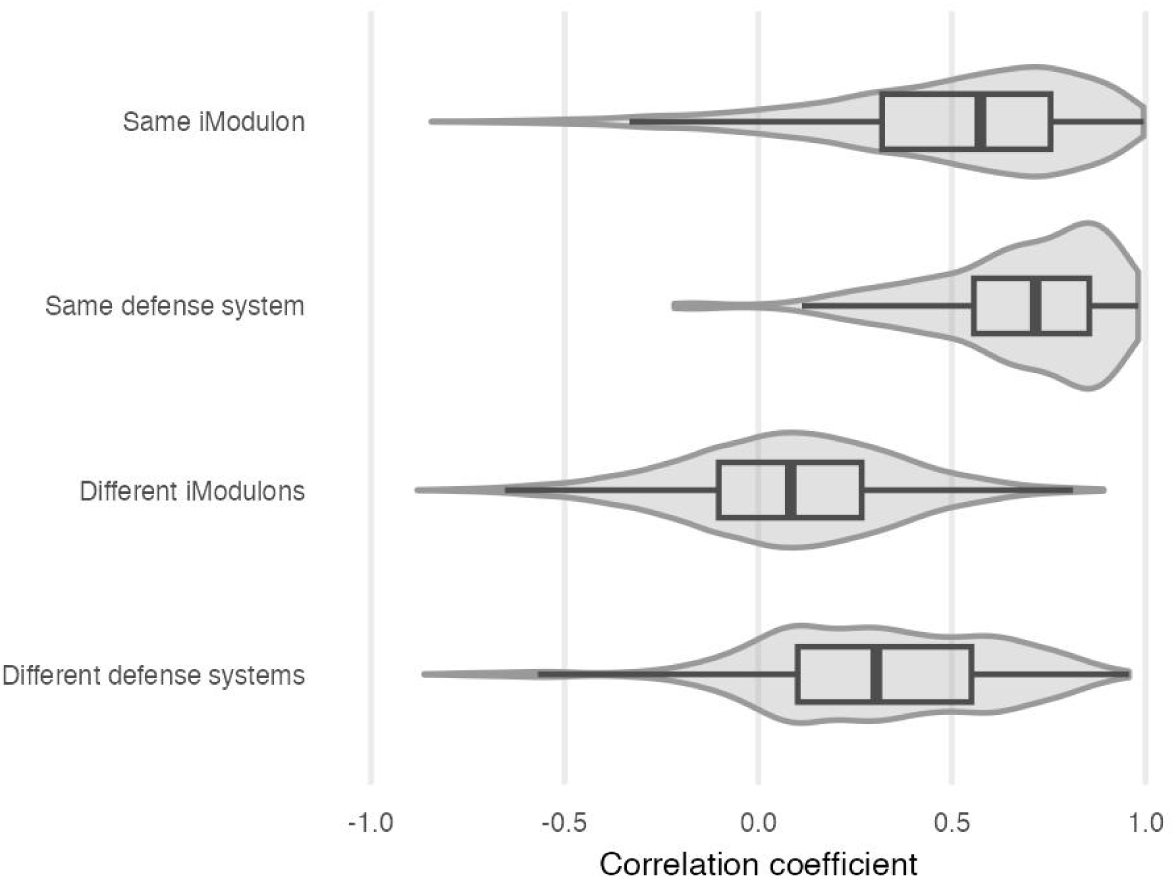
Correlations between defense systems and iModulons. Correlation of transcription patterns between genes of the same and different defense systems across the 14 species of the iModulons database. Defense systems correlations are contextualized by displaying correlations between genes within and between iModulons^23^.

**Supplementary figure 6:**
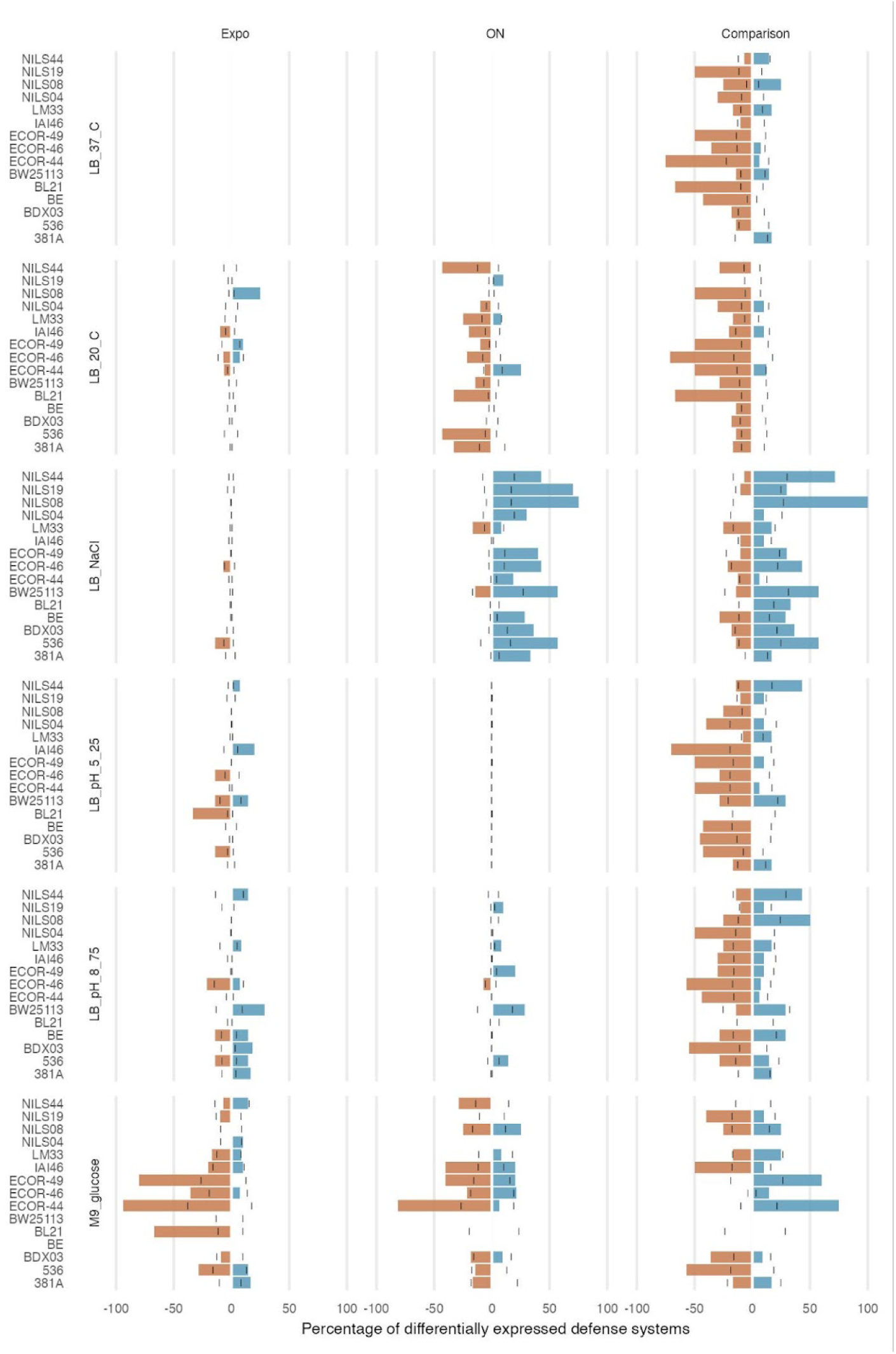
Strain-level differentially expressed defense systems across conditions. Similar to Figure 3G for all individual strains of the Picard-T dataset. Number of differentially up-or down-regulated defense systems compared to the average number of differentially expressed genes across conditions compared to the reference (Lennox LB 37°C, or condition-specific exponential phase).

**Supplementary figure 7:**
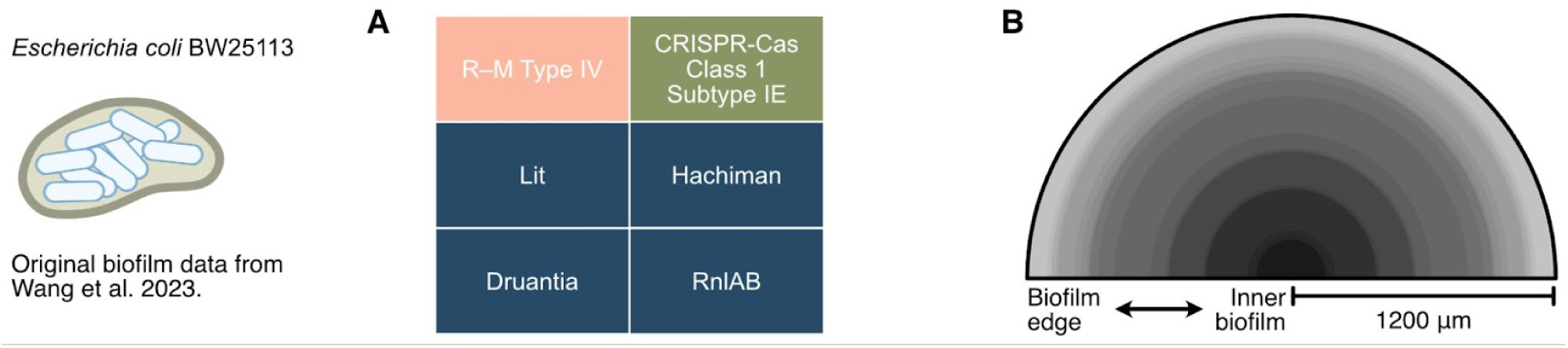
Spatial transcriptomics in a biofilm. A: Overview of the defense system encoded by the model *E. coli* strain from the spatial biofilm transcriptomic study. B: Schematic overview of the spatial resolution across the biofilm.

**Supplementary figure 8:**
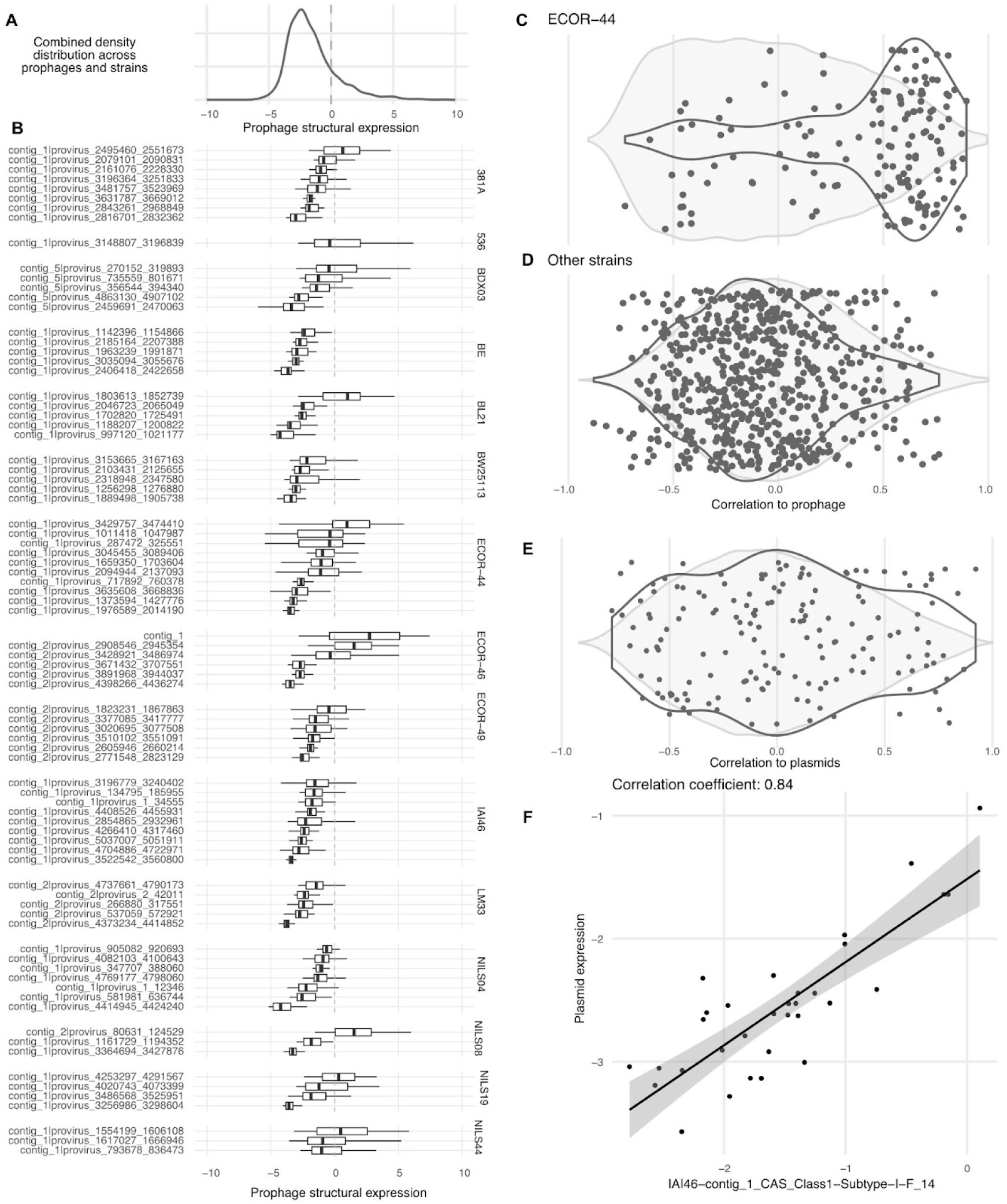
Expression of MGEs and defense systems. A-B: Distribution of prophage expression levels based on structural protein expression. C-D: The correlation between defense systems and prophages differs strongly between ECOR-44 and other strains. E-F: Correlations between plasmid and defense system expression.

**Supplementary figure 9:**
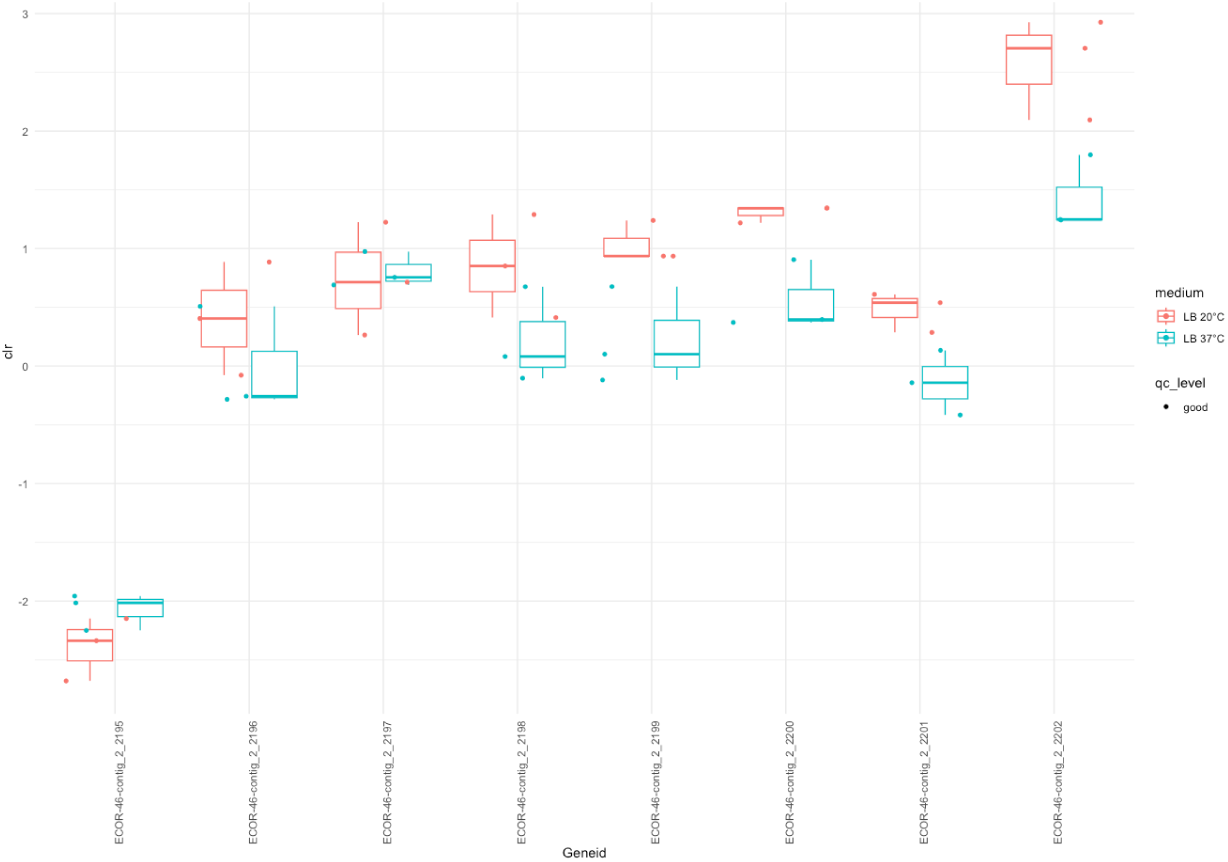
CRISPR-Cas expression in ECOR-46. Most genes of the CRISPR-Cas system of *E. coli* strain ECOR-46 are overexpressed in LB 20°C compared to 37°C.

**Supplementary figure 10:**
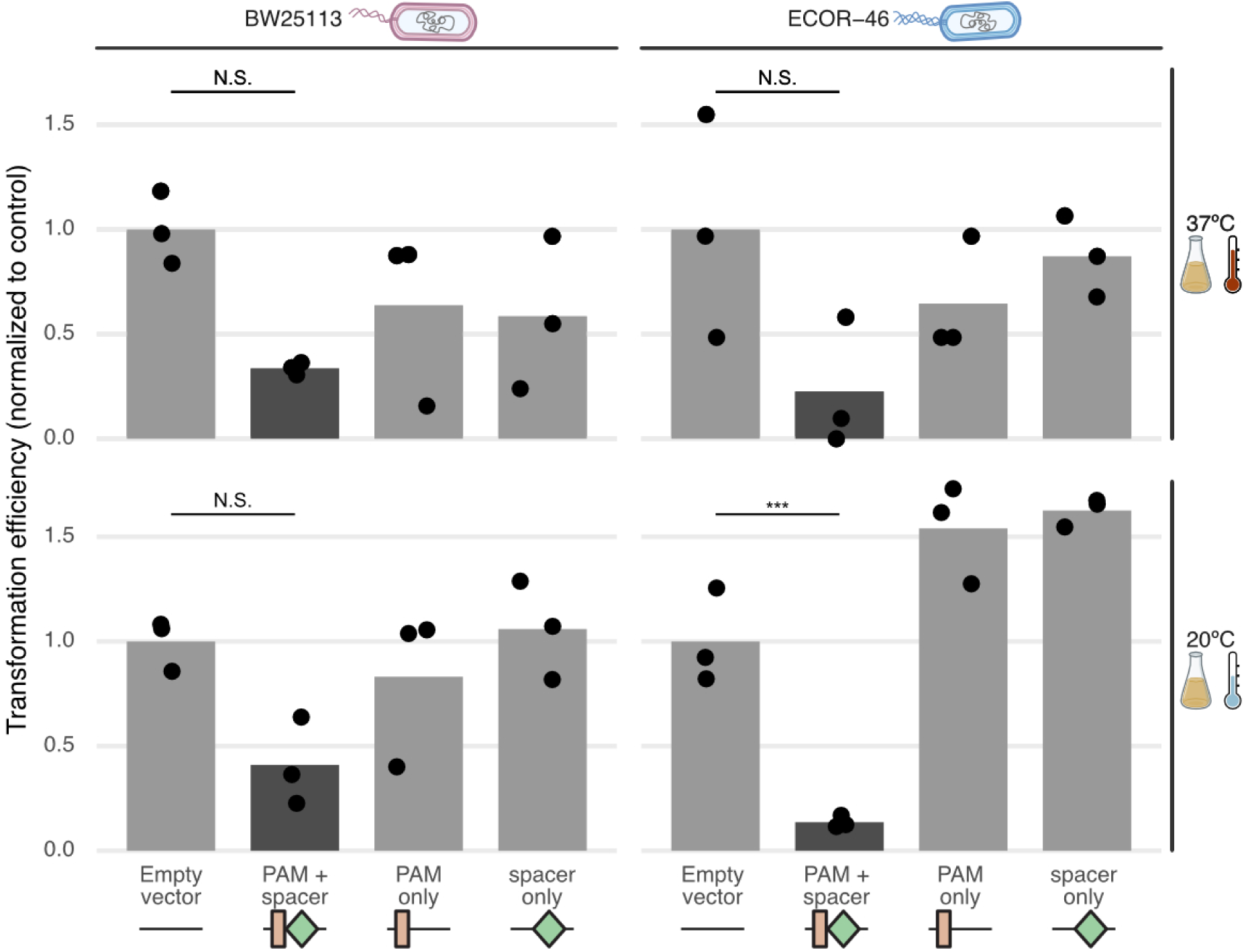
CRISPR-Cas functional validation across strains and conditions. Transformation efficiency across strains and conditions. Significance is based on within-strain and within-condition pairwise t-tests with FDR corrections. Results are only shown for the control vs PAM+spacer comparison for visualization purposes.

## Notes

### Competing Interest Statement

The authors have declared no competing interest.

